# An integral role for timing in interception

**DOI:** 10.1101/155531

**Authors:** Chia-Jung Chang, Mehrdad Jazayeri

## Abstract

Timing is critical for myriad behaviors in dynamic environments. For example, to intercept an object, the brain must compute a reliable estimate of time-to-contact (TTC). Prior work suggests that humans compute TTC using kinematic information such as distance and speed without explicitly relying on temporal cues, just as one would do in a physics classroom using kinematic equations. Considering the inherent uncertainty associated with estimates of speed and distance and the ability of human brain to combine different sources of information, we asked whether humans additionally rely on temporal cues. We found that humans actively integrate speed information with both explicit and implicit timing cues. Analysis of behavior in relation to a Bayesian model revealed that the additional temporal information helps subjects optimize their performance in the presence of measurement uncertainty. These findings suggest that brain’s timing mechanisms are actively engaged while interacting with dynamic stimuli.

## Introduction

Imagine intercepting a moving ball on a pool table as it bounces around hitting different edges. Does one only process kinematic information such as distance, position and speed, or does one additionally pay attention to *when* the ball hits the edges? At first glance, the answer seems trivial: since kinematic variables and time are directly related through kinematic equations (e.g., t = d/v), there would be no additional advantage in tracking time. What if internal estimates of speed and position were unreliable, for example, the lights were too dim to clearly see the ball? In that case, one may choose to pay attention to when the ball hits the edges to improve accuracy. This example highlights a general, important, and unresolved question in sensorimotor processing: do humans actively engage brain’s timing mechanisms when interacting with dynamic stimuli, or do they solely rely on kinematic information?

We aimed to address this question using a virtual object interception task. When intercepting a moving object, one has to estimate when the object reaches a desired target location, a variable that we will refer to as time-to-contact (TTC). Early studies hypothesized that TTC is derived from the rate of expansion of an object’s retinal image^1–3^. Later, it was suggested that TTC is derived indirectly from an object’s speed and position^4–13^. For example, when an object moves with a fixed speed, TTC would be computed by dividing perceived distance by perceived speed. However, as the example of the pool table illustrates, the inherent variability in the measurement and processing of kinematic information^14–20^ renders temporal cues highly relevant for the estimation of TTC. Humans’ ability to measure time intervals independent of kinematic cues is well-documented^21–24^. Furthermore, decades of research indicate that humans can efficiently combine multiple sources of information^14, 25–32^. Therefore, we hypothesized that humans integrate kinematic information with explicit and implicit temporal cues to derive better estimates of TTC (**Box 1**).

#### Box 1 Schematic paradigm and hypothesis for object interception.

(a) The overall logic of the experimental design. Subjects are asked to press a key when an bar moving with speed, *v*, would arrive at a target. The movement path is divided into a first section where the bar is visible, and a second section where the bar is invisible (i.e., virtually occluded). The subscript 1 and 2 are used to denote the distance (*d*), speed (*v*) and duration (*t*) of the two sections, respectively. Subjects’ behavior is evaluated by comparing the actual time-to-contact (*TTCa*) to the produced time-to-contact (*TTCp*), both of which are measured with respect to the moment the bar goes behind the occluder. (b) An estimate of TTC (denoted *TTCe*) can be derived by applying an appropriate transformation (noted as function, *f*) to measured stimulus parameters (*v_m_*, *t_n_*, *d_1m_*, *d_2m_*). The key question we focus on is whether subjects rely solely on the speed (*v_m_*), or they additionally incorporate temporal cues (*t_m_*).

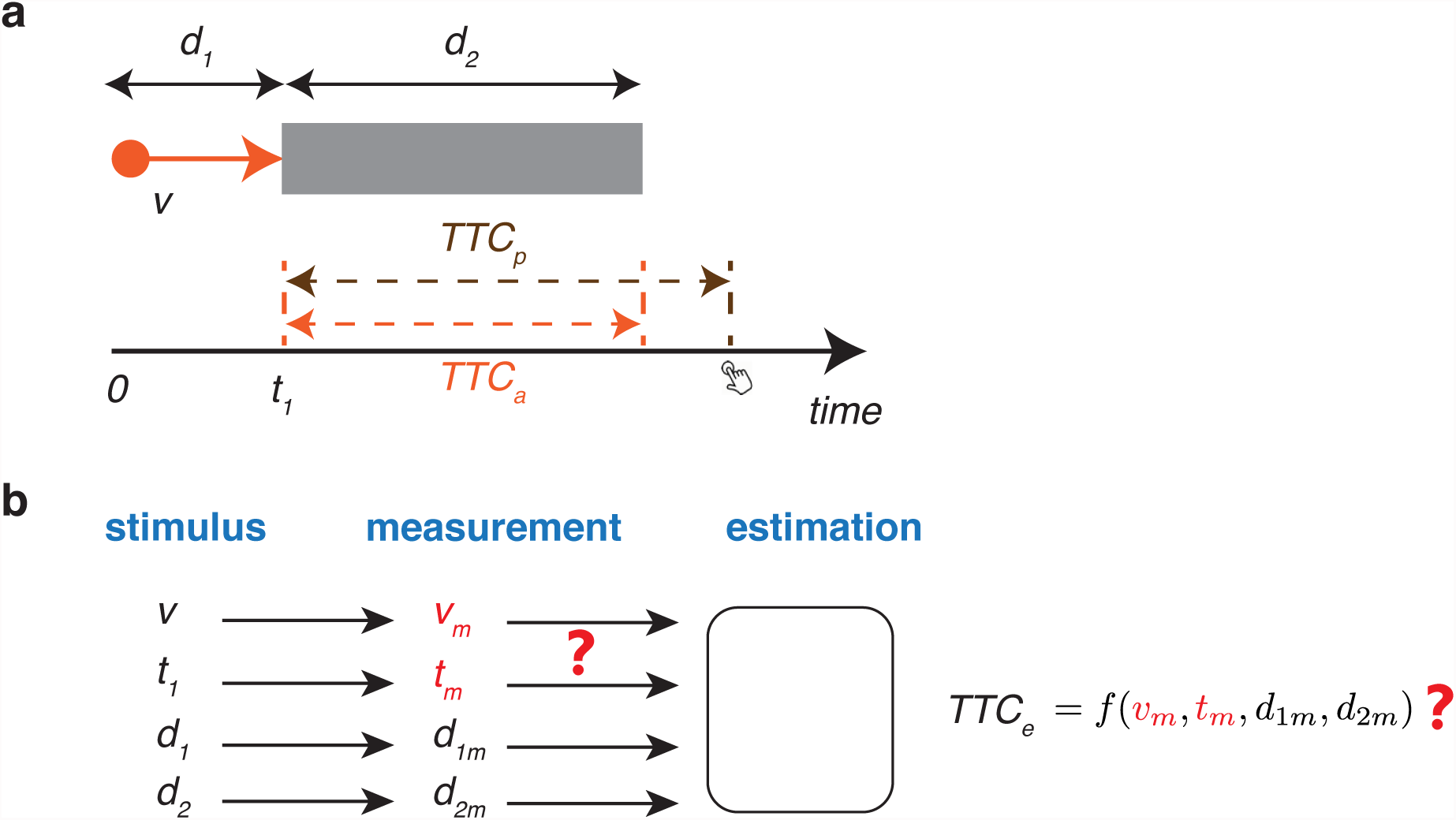

To test this hypothesis, we designed a series of experiments in which the subjects had to press a key when a bar moving along a linear path would arrive at a target position (**Fig. 1a**). While moving, the bar was sometimes visible and sometimes occluded. By varying the temporal structure between visible and occluded part of the path, we manipulated the reliability of the speed and the temporal information independently to compute TTC. Consistent with our hypothesis, we found that subjects actively used temporal information to improve their estimates of TTC.

**Figure 1.**
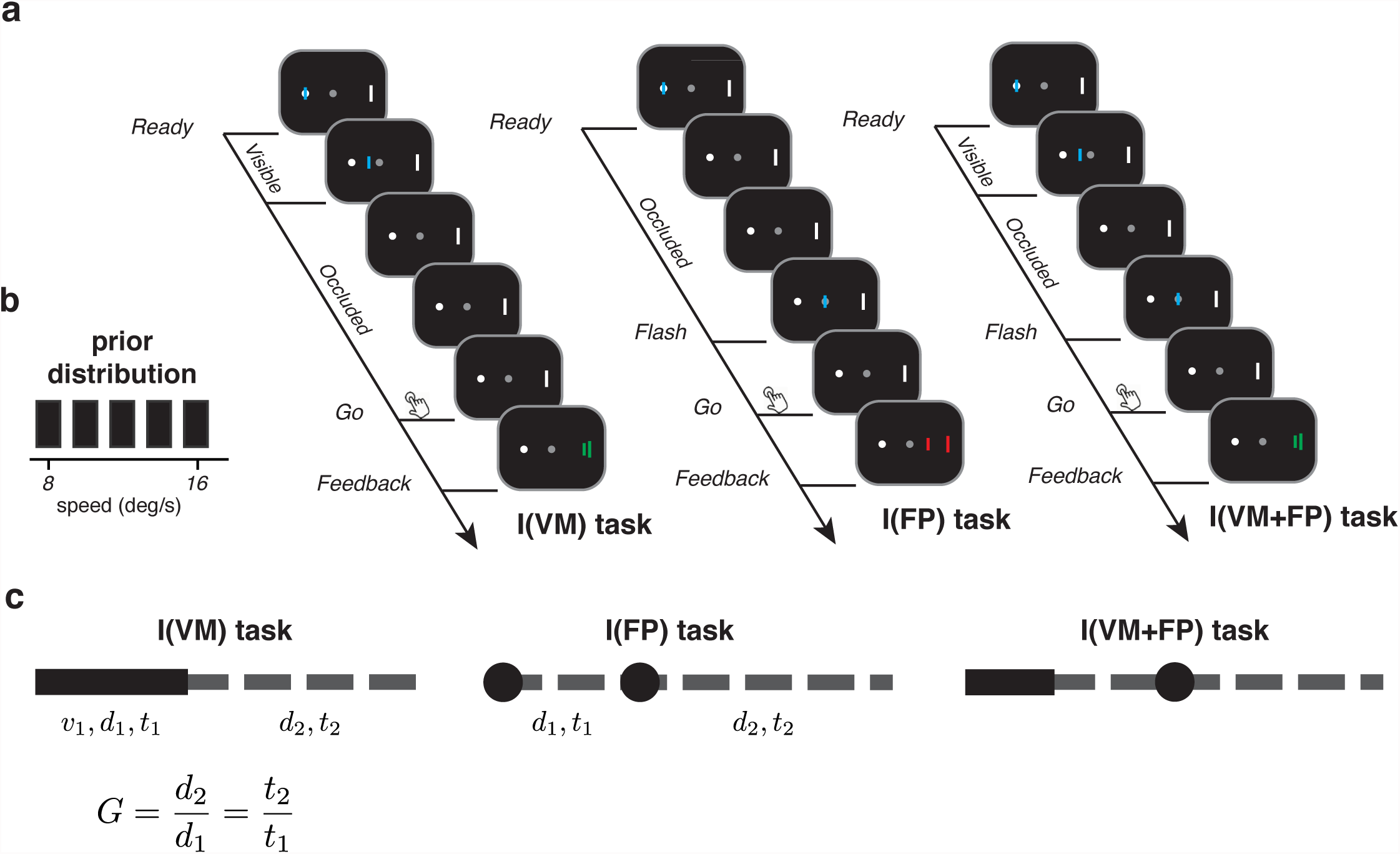
Experimental design and behavioral tasks. (**a**) Behavioral tasks. A bar moved from an initial point to the left of the fixation point to a target point to the right of the fixation point. The initial and target points were present throughout the trial. Subjects had to press a key when they judged the moving bar to have arrived at the target. Using this basic design, we tested subjects behavior in three conditions. In the I(VM) task (Interception with Visual Motion), the stimulus movement was initially visible and then invisible as if behind an (imaginary) occluder. In the I(FP) task (Interception with Flashed Position), the motion was not displayed and the stimulus was only flashed at the starting point and when it reached the fixation point. In the I(VM+FP) task, both the initial movement and the intermediate flash at the fixation point were displayed. In all trials, we provided feedback by presenting the position of the stimulus at the time of keypress. To reinforce accuracy, the target position and the stimulus feedback were shown in green when *TTCp* was within an experimentally defined window around *TTCa*, and red otherwise (see Methods). (**b**) Prior distribution of the stimulus speed. Speed was sampled from a discrete uniform distribution from 8 to 16 degree per second, and varied across trials. (**c**) Simplified symbolic representation of I(VM), I(FP), and I(VM+FP) conditions. The circles correspond to times when the stimulus was flashed, the solid lines to when the motion was displayed, and the dashed lines to when the stimulus was occluded. We used this symbolic representation as key for other figures.

To better understand the nature of the underlying computations, we compared subjects’ behavior to that of an ideal Bayesian observer who optimally integrates speed and timing information. Similar to work in other sensorimotor domains^14^, ^25–32^, the model was able to accurately capture subjects’ estimation strategy indicating that humans efficiently integrate prior statistics with measurements of both speed and elapsed time. These results highlight a hitherto unappreciated function of the brain’s capacity to utilize time – independent of speed and distance – to inform sensorimotor function while interacting with dynamic stimuli.

## Results

### Experiment 1: Interception performance benefits from explicit timing cues

We first asked whether humans are capable of integrating speed information with temporal cues to improve their estimate of TTC in an interception task. To do so, we asked subjects to intercept a moving bar in three conditions: one with a speed cue, one with a timing cue, and one with both cues present (**Fig. 2a**). In the first condition, the bar was visible only in the early part of the path, and then was occluded before it reached the target position. Subjects had to measure the speed of the bar from an early visible segment and use that to compute when the bar would reach the target position at the end of the occluded segment. We denote this condition by I(VM) as a shorthand for Interception in the presence of Visible Motion.

In the second condition, the motion was invisible but the position of the bar was flashed at the beginning of the path and when it reached the central fixation point, which was halfway along the path. TTC had to be computed based on the interval between two flashes along the path. We placed the timing cue around the fixation point to avoid causing a gaze shift in response to the flash, which could impact the subjects’ overall estimation strategy. We denote this condition by I(FP) as a shorthand for Interception in the presence of Flashed Position.

**Figure 2.**
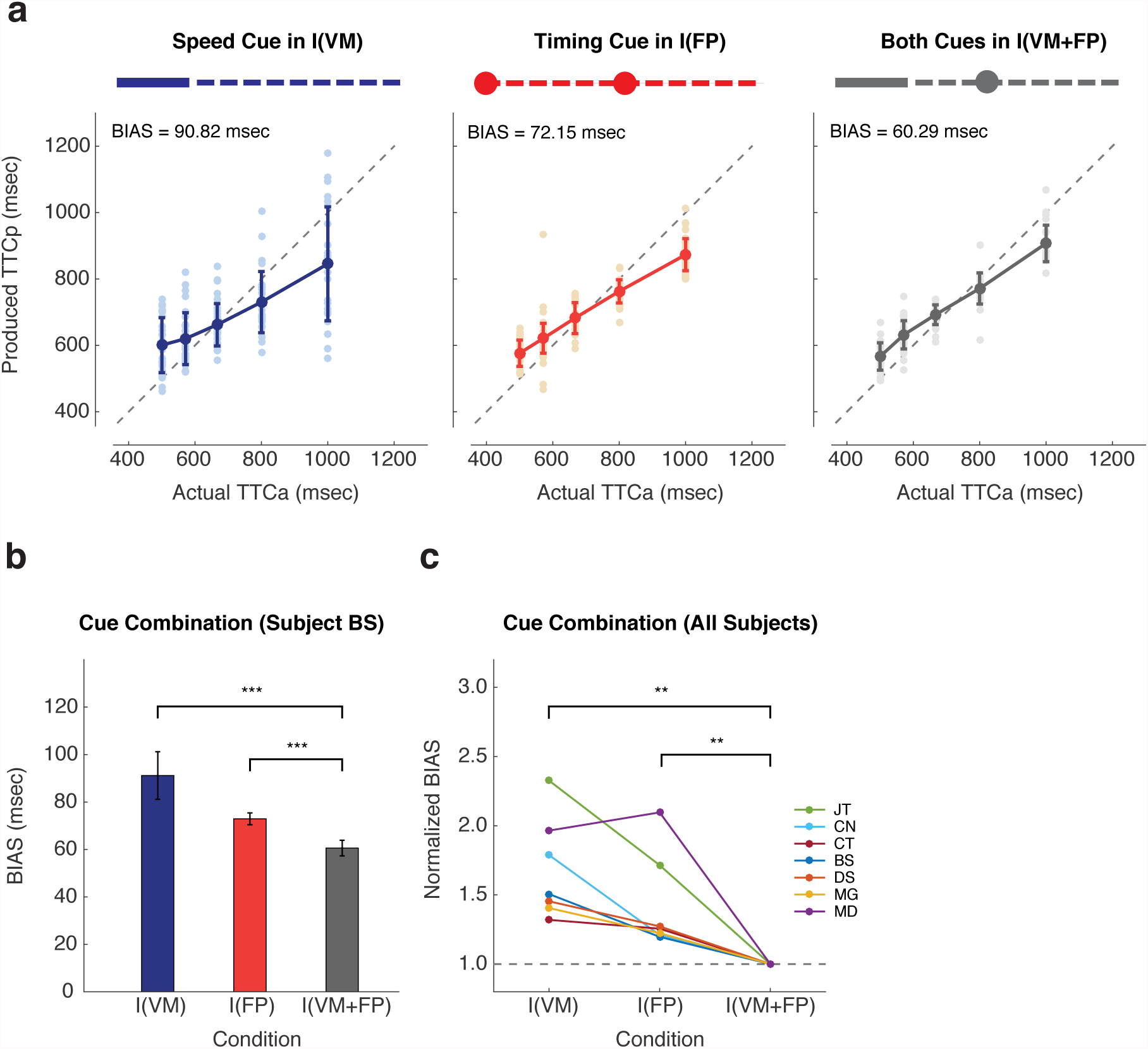
Interception using speed and explicit timing cues (Experiment 1). (**a**) Behavior of a typical subject for different conditions in Experiment 1. The left panel corresponds to the I(VM) task were the speed of the stimulus was evident from the initial visible section of the movement. The middle panel corresponds to the I(FP) task in which the stimulus was flashed at the starting point and the central fixation point. The right panel corresponds to the I(VM+FP) task where both the initial speed and the intermediate flash were presented. Performance was quantified by comparing subject’s produced time-to-contact (*TTCp*) to the actual timeto-contact (*TTCa*). *TTCa* was defined as the time between when the bar reached the central fixation to when it reached the target. *TTCp* was defined as the time between when the bar reached the central fixation to when the button was pressed. Light dots and dark circles show *TTCp* in each trial and the corresponding averages for each *TTCa*. BIAS in each plot was quantified as the average error over the five distinct *TTCa* of the prior distribution, i.e., the root mean square of differences between five solid dark circle and the corresponding diagonal dash line on the plot. (**b**) BIAS comparison across conditions for a typical subject. We estimated the standard error through resampling data with 100 repetitions. BIAS was smaller for the I(FP) compared to I(VM), and smallest in the I(VM+FP) condition. (**c**) Normalized BIAS across conditions for all subjects (N = 7) shown in different colors. Normalized BIAS was obtained by dividing BIAS in all conditions with BIAS in the I(VM+FP) condition. Across subjects, BIAS patterns were similar to the typical subject in panel b.

In the third condition, the bar was visible early on, and additionally, its position was flashed briefly at central fixation point, i.e., halfway between the initial and target location, giving subjects the opportunity to measure both speed and timing information. We denote this condition by I(VM+FP).

The actual time-to-contact (*TTCa*) was defined as the time from when the bar passed the central fixation to when it reached the target. We compared *TTCa* to the interval between when the bar passed the central fixation and subjects pressed the button. We refer to this interval as the produced time-to-contact (*TTCp*). For I(FP) and I(VM+FP) conditions, *TTCp* was straightforwardly computed from the time when the bar was flashed to when the button was pressed. For the I(VM) condition, because the bar was not visible throughout the path (not flashed at the fixation point), we estimated *TTCp* by appropriately scaling the response time by the occluded segment.

As evident from the *TTCp* pattern for a typical subject (**Fig. 2a**), subjects were able to perform the task in all three conditions with different degrees of of sensitivity. *TTCp* values were variable and systematically biased toward the mean. We quantified this regression to the mean by computing a BIAS term that quantifies the overall deviation from the identity line (see Methods; **Fig. 2b**). The BIAS was significantly smaller when both cues were available compared to the speed cue condition (t198 = 26.6435, p < 0.001, hedges’ g = 3.7537) as well to the timing cue condition (t198 = 27.4602, p < 0.001, hedges’ g = 3.8687). This reduction in BIAS was observed for all the subjects (**Fig. 2c**) and was significant across subjects

(Wilcoxon one-side signed-rank test, statistics = 28, p < 0.01), suggesting that humans are capable of integrating speed information with temporal cues to reduce uncertainty.

While this result is consistent with subjects integrating the two cues, it is also possible that the subjects did not integrate the two cues and instead used the timing cue (flash at the fixation point) to simply reset their subjective estimate of the position of the bar to the central fixation. To test this possibility, we tested a subset of subjects in a cue conflict paradigm in which the flash at the central fixation (FP) was jittered by –100, 0, or 100 msec relative to when the bar reached the central fixation (**Supplementary Fig. 1a**). To understand the logic of this experiment, let us predict subjects’ estimated time-to-contact (*TTCe*) under various hypotheses for a case where the flash is presented 100 ms later that when the bar reaches the fixation point. If a subject only relies on the timing cue, they would overestimate the time from motion onset to the flash by 100 msec; this would predict that *TTCe* would be 100 msec longer than *TTCa*. If a subject only relies on speed, then the lagging flash would not influence the subject’s behavior. However, since we quantify *TTCp* with respect to the time when the bar is flashed, we would register a *TTCe* that would be 100 shorter than *TTCa*. As a third hypothesis, let us consider that a subject would use the flash to reset the position, and use the speed to estimate *TTCe*. For this hypothesis, the *TTCe* would remain the same as when the subject only used the speed cue. Finally, if the subjects were to integrate the two cues, we would see a bias in *TTCe* in the direction of the jitter that would be less than 100 msec away from the non jittered condition. The results were consistent with the last hypothesis of integration and could not be explained by hypotheses including the position-reset hypothesis (**Supplementary Fig. 1a**).

### Experiment 2: Interception performance benefits from inherent timing cues

Experiment 1 demonstrated that humans were able to integrate timing cue with speed information. However, this could have been due to the presentation of an explicit timing cue (i.e., the flash a the fixation point). Therefore, we asked whether subjects utilize the timing information from the visible portion of the motion even if no explicit flash at the fixation point is presented. To validate the role of time as an additional cue, it was important to make sure that longer visible motion did not additionally improve subjects’ estimate of the speed. Therefore, as a first step, we measured the ability of subjects to estimate speed during interception with different visible lengths ranging from 0.625 to 5 degree in log scale while keeping the occluded distance fixed (**Supplementary Fig. 2**). We evaluated performance by measuring subjects’ root mean squared error (RMSE). We found that performance improved significantly as the visible lengths increased from 0.625 to 1.25 degree (paired-sample t-test, t_399_ = 56.61, p < 0.001) and saturated afterwards (paired-sample t-test, t399 = 0.9031, p = 0.3670). In other words, the fidelity of the speed estimate saturated at a visible length of 1.25 degree.

We then conducted an interception task where we evaluated the relevance of the timing cue by changing the visible length compared to the occluded length. Importantly, in all conditions, the visible length was beyond the saturation point in the I(VM) task. This ensured that any improvement in performance was not due to an improvement of speed estimates. We tested subjects’ performance in three conditions. In all conditions, the occluded length was fixed (d2 = 8 degree). Across conditions, the ratio of the occluded length (d2) to the visible length (d1) was varied by a gain factor (G = d2/d1). The three gain factors were 0.667, 1, and 1.6.

**Figure 3a and 3b** shows the performance of a typical subject in the three conditions. Surprisingly, the best performance was not associated with G = 0.667 when the visible length was longest. Instead, RMSE was smallest when the visible and occluded lengths were equal (G = 0.667, t_198_ = 20.3981, p < 0.001, hedges’ g = 2.9308; G = 1.6, t_198_ = 22.9261, p < 0.001, hedges’ g = 3.2299), which we refer to as the identity condition. The same was true across subjects (**Fig. 3c**; Wilcoxon one-side signed-rank test, statistics = 28, p < 0.01) revealing a systematic and consistent improvement of performance in the identity condition.

**Figure 3.**
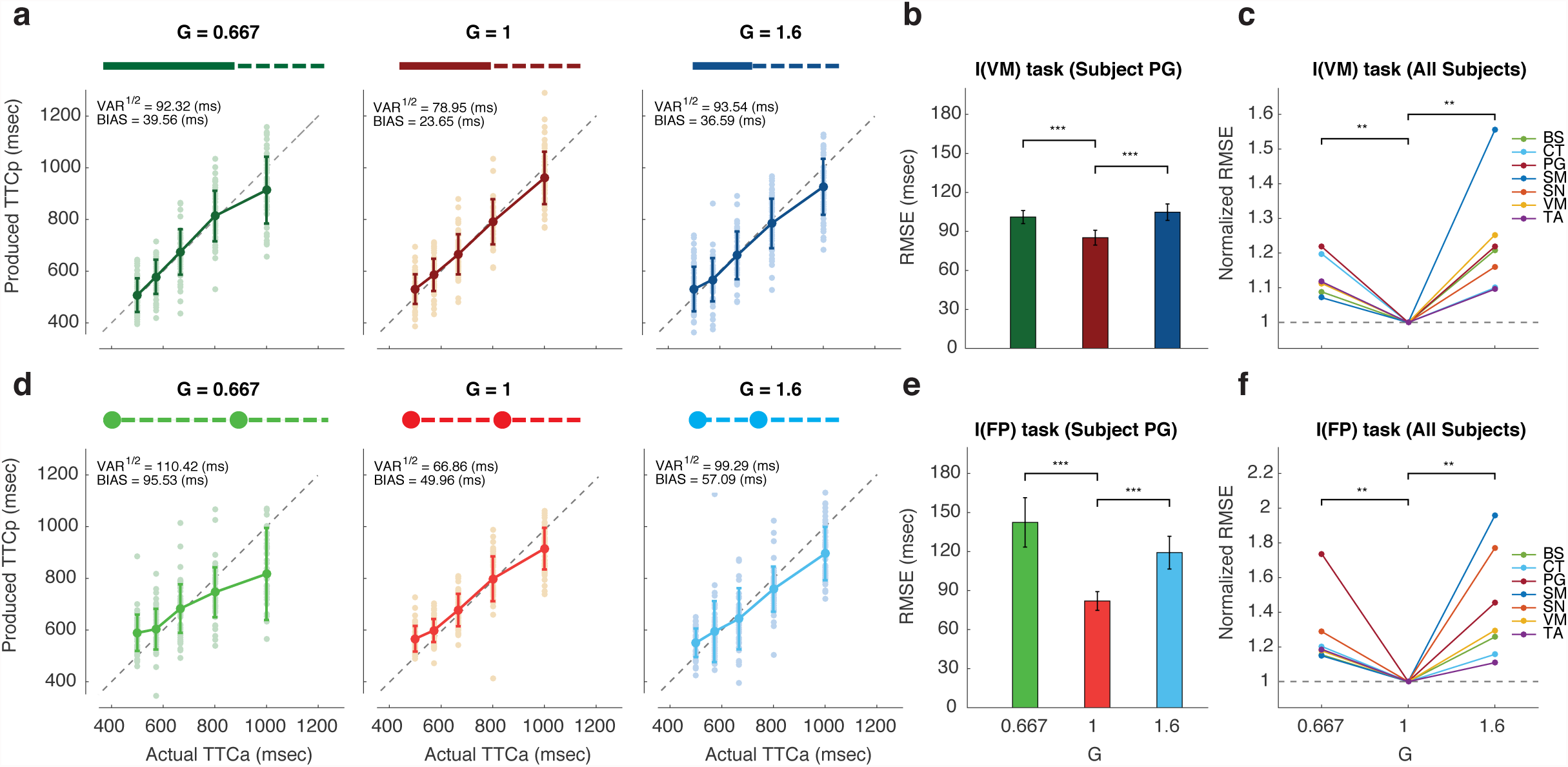
Interception using speed and implicit knowledge of temporal context (Experiment 2) (**a**) Behavior of a typical subject in three variants of the I(VM) task with three different visible lengths and the same occluded length. Each condition was identified by a gain factor (G) that quantified the ratio of the occluded to visible length. Since the bar moved at a constant speed throughout each trial, the gain also reflected the ratio of the duration of occluded and visible sections of the path. Performance was quantified by comparing subjects’ produced time-to-contact (*TTCp*) to the actual time-to-contact (*TTCa*). Light dots and dark circles show *TTCp* in each trial and the corresponding averages for each *TTCa*. BIAS was defined as described in Figure 2. VAR is the average variance of *TTCp* over the five intervals tested (*TTCa*) of the prior distribution. (**b**) Comparison of performance in terms of RMSE across conditions for a typical subject in I(VM) task. We estimated the standard error of RMSE through resampling data with 100 repetitions. (**c**) Normalized RMSE as a function of G for I(VM) task across all subjects (N = 7). RMSE in each condition was divided by RMSE when the gain was identical (G = 1). Different colors lines correspond to different subjects. (**d**) Behavior of a typical subject for I(FP) task with different gains (G) in Experiment 2. (**e**) Comparison across conditions for a typical subject in I(FP) task. (**f**) Normalized RMSE as a function of G for I(FP) task across all subjects (N = 7). For all subjects and in both tasks, RMSE was smallest for G=1 indicating best performance when the visible and occluded lengths were the same length (identity context, see main text).

The same group of subjects were also tested in the I(FP) task, and for the same three gains. As evident from the behavior of the same typical subject, RMSE was smaller when the measurement and production intervals were the same (**Fig. 3d and 3e**) compared to when the measurement period was longer (G = 0.667, t_198_ = 29.7316, p < 0.001, hedges’ g = 4.1887), or shorter (G = 1.6, t_198_ = 25.6390, p < 0.001, hedges’ g = 3.6122). This effect was present across subjects (**Fig. 3f**; Wilcoxon one-side signed-rank test, statistics = 28, p < 0.01) indicating that spatiotemporal identity helped subjects improve their estimate of TTC. We also compared subjects’ performance in the identity condition between the I(FP) and I(VM) conditions (**Supplementary Fig. 3a**). RMSE was consistently and significantly smaller in the I(VM) condition (Wilcoxon one-side signed-rank test, statistics = 3, p < 0.001). This ruled out the possibility that subjects switched to a pure timing strategy in the identity context. These results suggest that subjects exploited the temporal structure to improve their performance.

### Experiment 3: Interception performance improves with temporal-identity context

Experiment 2 clearly demonstrated that interception was most accurate in the identity context when the visible and occluded parts of the path were identical. This is consistent with our hypothesis that performance benefited from the fact that the visible and occluded intervals had the same duration (i.e., temporal identity) allowing subjects to more accurately estimate TTC. However, it is also possible that this improvement was because the visible and occluded parts had the same length (i.e., distance identity) allowing subjects to better estimate distance. The latter hypothesis seems unlikely given that the occluded distance was fixed throughout all experiments. Nonetheless, we conducted an additional experiment to assess the relevance of temporal versus distance identity in interception performance.

Since distance and duration are related through speed, the only way to dissociate the two is to make the speed of the bar differ between the visible and occluded parts of the path. Therefore, we designed a variant of the interception task in which, unbeknownst to the subjects, the speed of the bar behind the occluder was made 1.25 times faster than the speed in the visible portion (**Fig. 4a**). The non-identical speed ratio enables us to create conditions in which the distance and temporal identity were dissociated. In one condition, the visible and occluded distances were the same creating distance identity (G_d_ = 1) without temporal identity (G_t_ = 1/1.25). In another condition, we matched the ratio of the distances to the ratio of the speeds (G_d_ = 1.25) to create temporal identity (G_t_ = 1).

**Figure 4.**
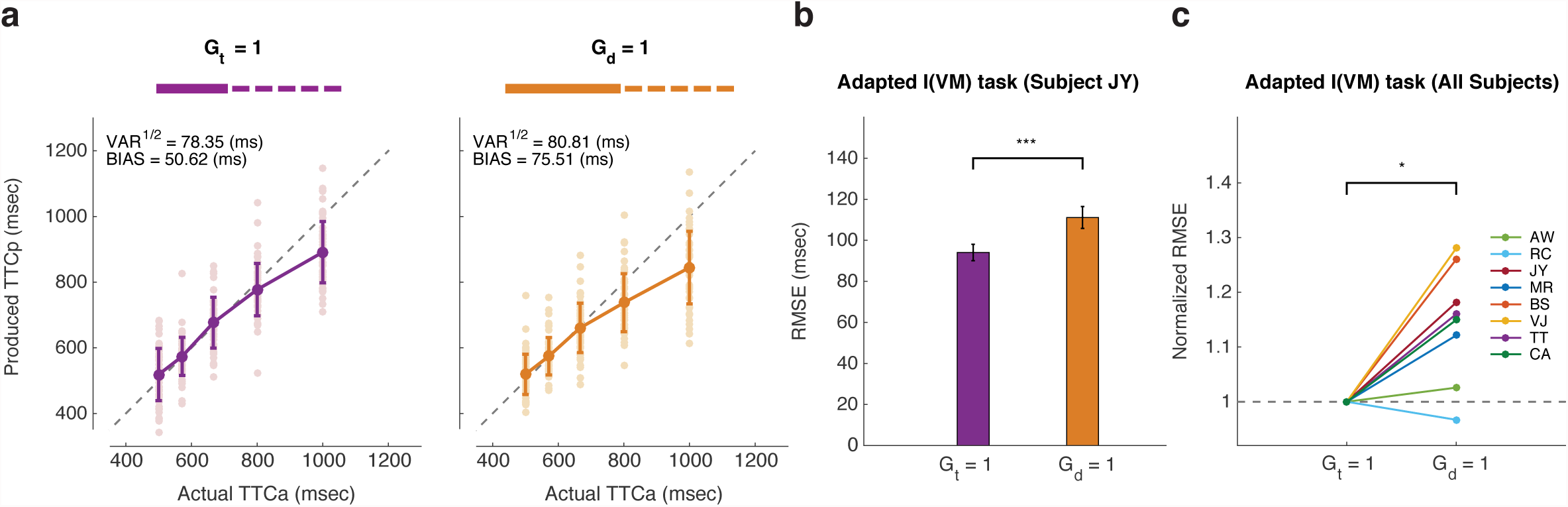
Interception using distance and temporal identity contexts (Experiment 3) (**a**) Behavior of a typical subject for two variants of the I(VM) task, the temporal identity context (*G_t_ = 1*) and the distance identity context (*G_d_= 1*). In both variants, unbeknownst to the subject, the speed behind the occluder was multiplied by 1.25 (25% faster than the visible section). *G_t_ = 1*: The durations of movement in the visible and occluded sections were the same. Because of speed difference between the two sections, the visible distance was shorter than the occluded distance. *G_d_ = 1*: The visible distance was same as the occluded distance, but the corresponding durations were different. (**b**) Comparison between the two conditions of I(VM) shown for a typical subject. We estimated the standard error of RMSE through resampling data with 100 repetitions. (**c**) Normalized RMSE across all subjects (N = 8). Different colored lines represented different subjects.

A new set of subjects was recruited for this experiment to ensure that any sensitivity to temporal context was not because of participation in previous experiments. Since subjects were not aware of the speed change behind the occluder, they could only adjust their performance based on feedback. We compared subjects’ performance between the G_d_ = 1 and G_t_ = 1 conditions. We reasoned that an observer that relies on the distance identity should have higher performance (lower RMSE) in the G_d_ = 1 condition. In contrast, an observer that relies on the temporal identity would have a lower RMSE in the G_t_ = 1 despite the fact that the distances between the visible and occluded parts are not the same.

We found that RMSE was lower for the temporal identity compared to distance identity condition as shown for a typical subject (**Fig. 4b and 4c**; t_198_ = 25.6431, p < 0.001, hedges’ g = 3.6127) and across subjects (Wilcoxon one-side signed-rank test, statistics = 34, p < 0.05). This finding further substantiates our conclusion that subjects rely more on temporal context to estimate TTC.

### Bayesian integration of speed and time explains interception performance

Experiments 1 to 3 established that subjects integrate speed and timing information to improve their performance. Another salient feature of subjects’ behavior across all conditions (**Fig. 2** and **Fig. 3** and **Fig. 4**), was that *TTCp* was biased toward the mean of the prior. This was true for the external timing cue tasks in Experiment 1, for the inherent timing tasks in Experiment 2, and in the control condition in Experiment 3, regardless of the ratio of the occluded to visible lengths (different values of G). Together, these observations suggest that subjects performance may be explained by a Bayesian model that integrates the prior with both the speed and timing information (**Fig. 5a**).

**Figure 5.**
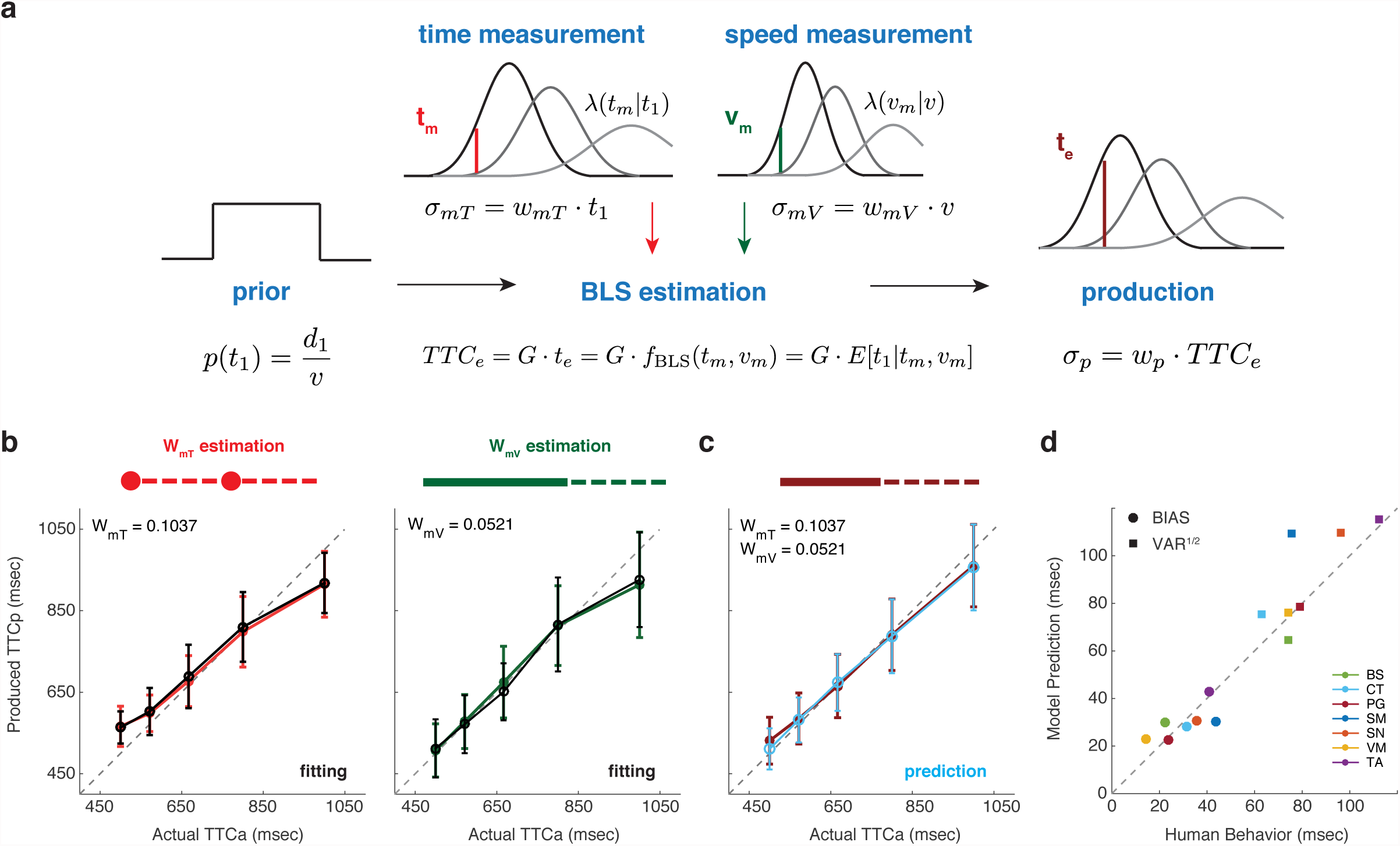
The Bayesian observer model of interception integrating speed and timing cues. (**a**) The Bayesian observer model for the I(VM) task. On each trial, the speed (*v*) was drawn from a uniform prior distribution. We used the relationship between the distance of the visible section (*d_1_*) and speed to express the prior in terms of the duration the bar is visible (p(*t_1_*); left). We assumed that the observer makes two conditionally independent measurements of *v* and *t_1_*, which we denoted by *v_m_* (red vertical line) and *t_m_* (green vertical line), respectively. We assumed that *v_m_* and *t_m_* are perturbed by zero-mean Gaussian noise with standard deviations (σ*_mV_* and σ*_mT_*_)_ proportional to *v* and *t_1_* (top Gaussian curves) with constant of proportionality of *w_mV_* and *w_mT_*, respectively. The Bayesian observer computes the posterior from the likelihood functions,λ(*v_m_*|*v*) and λ(*t_m_*|*t_1_*), and the prior, and uses a Bayes-Least-Squares (BLS) estimator, *f_BLS_*, to infer the movement duration in the visible section, which we denoted by *t_e_* (brown vertical line) from *v_m_* and *t_m_*. This estimate is then multiplied by the distance gain (G) to obtain an optimal estimate of time-to-contact (*TTCe*). Finally, the model incorporates motor variability via additional noise in the production stage. We modeled this noise as a sample from a zero-mean Gaussian with standard deviation scaling with *TTC_e_* with scaling factor *w_p_*. (**b**) The left panel _(_*w_mT_ estimation*) shows the behavior of a Bayesian observer model (red) fitted to the data (black) for a typical subject in I(FP) task with G = 1. Since the movement of the bar in the I(FP) task is not visible, we estimated w_mT_ from a Bayesian model that relies on the prior and *t_m_*, but not *v_m_*. The right panel (*w_mV_ estimation*) shows the Bayesian model (green lines) and the corresponding data (black) for the I(VM) task with G = 0.667. In the I(VM) condition, the observer has access to both speed and time. Therefore, we estimated *w_mV_* from a Bayesian model that uses the prior, *t_m_* and *v_m_* with *w_mT_* inferred from I(FP) with G = 0.667. (**c**) Behavior (black) and model prediction (blue) for a typical subject in the I(VM) task with G = 1. The prediction was made based on a Bayesian model whose *w_mT_* and *w_mV_* were derived in (**b**). (**d**) Comparison of summary statistics (BIAS and VAR^1/2^) between human behavior (abscissa) and predictions from a Bayesian model (ordinate) across subjects (N = 7). BIAS and VAR^1/2^ of the model was computed based on averages of 100 simulations of the Bayesian observer model. Different colors correspond to different subjects

To test this hypothesis rigorously, we developed an ideal Bayesian observer for the task. We assumed that the ideal observer made two conditionally independent measurements while the moving bar was visible, one associated with the speed of the bar (*v_m_*), and another associated with the duration of the visible interval (*t_m_*). Following previous work, we assumed that these measurement were subject to scalar variability^26, 33–36^. In particular, we assumed that the standard deviation of noise on speed scaled with the bar’s speed (*v*) with constant of proportionality (*w_mV_*) and standard deviation of noise on elapsed time scaled with visible duration (*t_1_*) with constant of proportionality (*w_mT_*). The ideal observer integrated the prior, *p*(*t_1_*), with the likelihood of the bar speed, λ(*v_m_|v*) and the likelihood of the visible duration, λ(*t_m_|t_1_*), and computed TTCe from the mean of the posterior. Since this observer minimizes the least-squares error, we will refer to this as the Bayes least-squares (BLS) estimator. To compare the model to subjects’ behavior, we augmented the ideal observer with a production stage by adding scalar noise with constant of proportionality (*w_p_*) to *TTCe* to values of *TTCp* that incorporated motor variability.

We first estimated *w_mV_*, *w_mT_* for each subject. In most Bayesian models, the model is evaluated by assessing the quality of model fits to the data. A more powerful approach is to fit the model to a training dataset and examine how well it explains a test dataset. An even more powerful approach is to fit the model to one set of conditions and ask whether it predicts data in another condition to which it was not fitted. We employed the last approach. For each subject, we estimated *w_mT_* from the I(FP) task with G = 1 (**Fig. 5b, left**), and *w_mV_* from I(VM+FP) in G = 0.667 (**Fig. 5b, right**), and used those estimates to predict subjects’ behavior in the I(VM+FP) in G = 1 (**Fig. 5c**).

To estimate *w_mT_*, we developed a Bayesian observer for the I(FP) task with G = 1. In this task, the sensory information provided was the interval between when the bar started to move and when it reached halfway along the path (over the fixation point), which we denote by *t_1_*. We fitted subjects’ behavior by a BLS estimator that only relied on the likelihood of *t1*, λ(*t_m_|t_1_*) and the prior distribution, *p*(*t_1_*). As shown for one subject (**Fig. 5b, left**) and consistent with previous work in a similar task^21^, ^37–39^ the model accurately captured subjects’ behavior.

Next, we estimated *w_mV_* from fits of the Bayesian model to the I(VM+FP) task when G = 0.667. For this fitting procedure, we used the corresponding *w_mT_* from the I(FP) task with G = 0.667 (see **Methods**). As shown for the same subject (**Fig. 5b, left**), the model successfully accounted for the behavior. Recall that in the I(VM+FP) task, we had made the visible length long enough so that subjects’ estimate of speed had saturated and was thus no longer dependent on G (**Supplementary Fig. 2**). This allowed us to safely use the fit to *w_mV_* derived from the G = 0.667 condition to predict behavior in the G = 1 condition.

Finally, we use each subject’s fits to *w_mV_*, *w_mT_* to predict the behavior in I(VM+FP) task when gain is one (G = 1). The model was able to predict the observed *TTCp* values as shown for one example subject (**Fig. 5c**) and captured the data’s summary statistics (BIAS and VAR) across subjects (**Fig. 5d**). This is remarkable considering that both *w_mV_* and *w_mT_* were estimated from other tasks, and provides strong support that subjects integrate prior information, speed information, and timing information to optimize their estimate of TTC.

To further evaluate the success of the Bayesian model in explaining how subjects integrate speed and timing information, we tested the model in Experiment 1 where the timing cue was provided explicitly by a flash at the fixation point. To create a predictive model, we used the same procedure as we did to predict behavior in Experiment 2. We estimated *w_mV_* from data in I(VM) condition, and *w_mT_* from data in I(FP) condition (**Supplementary Fig. 3b**), and used those values to predict behavior in I(VM+FP) condition (**Supplementary Fig. 1b**). Again, the model successfully captured the statistics of subjects’ behavior suggesting that the brain is optimized for integrating speed and timing information during interception tasks regardless of how timing information is provided.

## Discussion

Our work builds on a large body of work investigating the computational principles of object interception. Early studies hypothesized that humans rely on variables derived from an object’s visual angle and its rate of expansion on the retina, of which the so-called tau is a classic example^1–3^. Later, this proposal was deemed inadequate as it failed to capture many empirical observations^4, 9, 40–42^. Most current models are based on the idea that interception relies on measurements of kinematic variables^6^, ^7^, ^11^, ^12^, ^43^, such as speed^6^, ^32^, ^44^, distance and/or depth^45^. This idea has also been used in experiments similar to ours where the object moves behind an occluder^6^, ^9^, ^46–48^. In those cases, it is assumed that humans estimate speed while the object is visible and use that estimate to predict future position of the object behind the occluder. This focus on kinematics is natural as it matches our intuition about the physics of how objects move. However, the algorithms the brain uses for object interception need not match our physics intuition. Here, we asked whether humans solely rely on kinematics (e.g., speed and distance), or do they additionally rely on temporal cues and contexts.

Real world object interception involves a decision to initiate a movement followed by online adjustments of the movement based on sensorimotor feedback. Although successful interception requires a tight coordination between the initiation and the subsequent adjustments, the two processes typically involve different computations^49^. The decision of when to initiate is, by and large, determined by a prediction of how long it would take to reach the object – i.e., time-to-contact (TTC), whereas the subsequent adjustment involves fine adjustments after the movement has been initiated. Here, we focused on the former asking how the brain determines TTC. To do so, we designed a virtual interception task in which subjects “intercepted” a moving bar by pressing a button when the bar reached a target position behind an occluder. With this design, we effectively eliminated the need for post-initiation adjustments. Our main objective was to investigate whether TTC was computed solely from estimates of bar kinematics (e.g., speed and distance), or whether subjects additionally relied on temporal cues and contexts.

We tested this question in two complementary sets of experiments. In the first set, we presented a brief flash showing the position of the bar after its disappearance behind the occluder. This flash provided explicit information about position and time of the bar but not its speed. Therefore, any improvement in performance due to the flash must be taking advantage of timing mechanisms in the brain. Results confirmed that subjects could intercept the bar without any speed information, and when the flash was presented along with additional speed information, subjects were able to integrate the two to improve their performance. This result complements a large body of evidence that humans are able to fuse sensory information from multiple modalities while making perceptual inferences^29^, ^31^, ^50^. `Note that the integration of speed and time is distinct from the indirect role that time would play by improving one’s estimate of speed^51–53^. As we demonstrated in a supporting experiment (**Supplementary Fig. 2**), the improvement of speed estimate with time saturates rapidly and cannot account for our finding. What our results reveal is that humans actively use elapsed time as an independent cue and integrate it with other visual cues when interacting with dynamic stimuli.

In the second set of experiments, we removed the explicit timing cue and instead asked whether subjects would naturally exploit implicit timing cues present in the temporal structure of the environment. To address this question, we designed an interception task in which we varied the interval the bar was visible. Based on recent work^54^, we reasoned that when the visible and occluded epochs have the same duration, subjects would automatically make use of this temporal identity to improve their performance. Subjects’ performance was remarkably improved in the temporal identity context compared to when the durations of the visible and occluded regions were not the same. Indeed, this experiment revealed a surprising aspect of human behavior: performance in the identity context was even better than when the occluded length was the same and the visible length was made longer. In other words, prolonging the visible portion was harmful to performance when it broke the temporal structure conferred by the identity context. This result powerfully demonstrated that the key factor driving the performance improvement was the presence of the identity context. This conclusion was reinforced by control experiments showing that the result was due to temporal – not distance – identity. Finally, we found that subjects’ ability to integrate speed and timing information reached performance levels similar to an ideal Bayesian observer, suggesting that human brain is inherently optimized to combine speed and time information for object interception.

These experiments lead to a simple and novel conclusion that humans actively engage timing mechanisms during interception. To put this finding in context, it is important to distinguish between the role of time during the visible and occluded regions of the path. When an object moves behind an occluder, subjects could no longer measure the object’s speed and thus have no choice but to rely on their sense of time. This idea was formalized by Tresillian and others in relation to human’s ability to extrapolate an object’s location behind an occluder^55^, ^56^. This is fundamentally different from what we propose; our findings indicate that humans actively integrate information about temporal contexts and events even when the object is visible. In other words, timing seem to be an integral component of how we interact with dynamics stimuli, both to better estimate where they are (when they are visible), and to infer where they might be (when they are occluded).

Our work does not address any potential role that timing information might play for the subsequent sensorimotor adjustments after movement initiation. It is possible that knowledge about temporal cues and contexts only inform movement initiation. This would indicate that temporal processing is only engaged during the cognitive and/or motor planning stage of object interception. This is consistent with numerous imaging and electrophysiological studies finding an important role for premotor and supplementary motor areas in timing^57–61^. Alternatively, knowledge about movement durations may also be used during movements although some studies have suggested that humans do not use timing information when they have access to movement related state-dependent information^62^, ^63^.

It is worthwhile considering why the role of time was not noted in prior research on object interception. We think that answer has to do with the simplicity of behavioral task used in laboratory settings (but see some studies using more naturalistic paradigms or done with virtual reality^47, 64^). Most experiments have not included rich spatiotemporal event and/or contexts, where temporal cues become relevant. However, real world examples of object interception take place in the presence of temporal statistics, spatial landmarks, and temporal events such as collisions and/or reflections, all of which make knowledge about time highly informative. A notable observation in our experiment was that subjects’ estimate of TTC was more accurate in the identity temporal context, possibly due to lower sensorimotor noise^54^. This improved sensitivity may be due to the fact that temporal identity creates a rhythmic structure between the relevant time intervals. If so, we would expect stronger effects when temporal events create sounds as auditory rhythms and/or integer ratios are constrained by strong internal priors^65^. For example, intercepting a bouncing ball may greatly benefit from the bounce sound, especially when visual information is uncertain (e.g., a dribbling a basketball without looking at the ball). These considerations highlight the need for future research to move beyond simple behavioral tasks and examine object interception in more naturalistic settings where the underlying dynamics are governed by richer spatiotemporal contexts. We speculate that doing so will further substantiate the importance of temporal events and contexts in processing dynamic stimuli.

## Online Methods

### Subjects

All subjects provided informed consent for experimental procedures which were approved by the Committee On the Use of Humans as Experimental Subjects at the Massachusetts Institute of Technology. Seven adult subjects participated in Experiment 1. A different group of seven adult subjects participated in Experiment 2. Another group of eight adult subjects participated in Experiment 3. All subjects had normal or corrected-to-normal vision.

### Procedures

Subjects sat in a dark, quiet room at a distance of approximately 50 cm from a display monitor with a refresh rate of 60 Hz and a resolution of 1920 by 1200 on an Apple Macintosh platform. Experiments were controlled by an open-source software (MWorks; http://mworks-project.org/). All stimuli were presented on a black background. Although eye movements were not monitored, all trials began with central fixation spot that subjects were asked to hold their gaze on the fixation point throughout every trial. Responses were made on a standard Apple Keyboard connected to the experimental machine.

We used three experiments to examine how people infer time-to-contact (TTC). Each experiment consisted of conditions whose order was randomized across subjects. Each condition was tested twice in two different days: the first session was used for training, and the second was used for the main test session but the first 25 trials were considered as warm-up and were excluded from the main analysis.

Subjects were asked to press a key when the bar reached the target position. Feedback was provided to indicate the actual bar position along the path when the key was pressed. The target position and the stimulus feedback were shown in green when the produced time-to-contact (*TTCp*) was within an experimentally defined window around the actual time-to-contact (*TTCa*), and red otherwise. To account for scalar variability, the window width was scaled with the actual time-to-contact (*TTCa*) with a constant of proportionality, *k*. The value of *k* was controlled by an adaptive one-up one-down procedure during training condition, and eventually reached a stable value, *k_0_*. We then set *k_0_* as feedback accuracy window in the main test session.

### Experiment 1

The objective of this experiment was to test whether subjects could improve their estimate of TTC by integrating motion and timing cues. The trials were structured as follows: subject pressed a key to initiate a trial. After a variable delay drawn randomly from a truncated exponential distribution (0.3-0.6 sec), a bar started moving horizontally from a starting point along a 16-degrees long linear path toward a target position at the end of the path (**Fig. 1a**). In each trial, the speed (v) of the bar was drawn from a discrete uniform distribution (**Fig. 1b**).

The experiment consisted of three different conditions in terms of the information subjects were provided with: one with motion cue, one with timing cue, and one with both (**Fig. 2a**). In the first condition, the interception path consisted of two sections: a section where the stimulus movement was visible and a section where it was occluded. The target was placed at the end of the occluded section. We denote this condition by I(VM) as a shorthand for Interception in the presence of Visible Motion. In the second condition, the motion was invisible throughout the path but the position of the stimulus was flashed at the beginning of the path and when it reached the central fixation point in the middle of the path. We denote this condition by I(FP) as a shorthand for Interception in the presence of Flashed Position. In the third condition, the stimulus movement was visible early on, and additionally its position was flashed when it reached the central fixation spot. Accordingly, we denote this condition by I(VM+FP). In tasks in which the position was flashed, the flashes lasted 100 msec. The distribution of sample interval (*t_1_*) between the start of the path and the time when the bar reached the fixation point was the same across the conditions. We also tested subjects in a cue conflict version of the I(VM+FP) in which the flash at the central fixation was jittered by –100, 0, or 100 msec relative to when the actual stimulus reached the central fixation (**Supplementary Fig. 1a**). These three jitter values were randomized and presented with equal probability.

### Experiment 2

The objective of this experiment was to test whether subjects could take advantage of temporal structure in the absence of an explicit temporal cue to improve their performance. The task was similar to the I(VM) condition in Experiment 1. A bar began to move from a starting point along a path. The bar was initially visible and then disappeared behind an occluder. Subjects pressed a key when the bar reached the target position at the end of the occluder. We tested subjects in three conditions (**Fig. 3a**). In all conditions, the distance between the fixation spot and target (d_2_) was set at 8 degrees while the visible length between the starting and the fixation spots (d_1_) was varied between 12, 8, and 5 degrees. We expressed these conditions in terms of the ratio of d_2_ over d_1_, which we define as a gain factor (G). The corresponding G for the three conditions were 0.667, 1, or 1.6. We recruited a new set of subjects for this experiment to make sure that participants were not made sensitive to timing cues due to prior experience with the I(FP) and/or I(VM+FP) tasks.

To evaluate the relative importance of speed and timing information, we also tested the newly recruited subjects with the same gain factors but in the I(FP) condition (**Fig. 3d**). However, all I(FP) conditions were tested after the subjects had completed the I(VM) conditions to avoid inadvertently sensitizing subjects to timing cues. Overall, Experiment 2 consisted of 6 conditions in total.

### Experiment 3

The objective of this experiment was to test whether the improved performance in Experiment 2 in relation to the identity context was related to the distance identity or temporal identity. To facilitate the description of these conditions, let us define the sample interval (t_1_) as the interval associated with the visible portion of the path, between the starting point and when the bar reached the central fixation (d_1_). Similarly, we define the target interval (t_2_) as the interval associated with the occluded part of the path (d_2_). Since the experiment involved changing the relative distances and/or durations, we additionally define two a distance ratio that corresponds to the ration of the occluded length to the visible length (G_d_ = d_2_/d_1_), and a duration ratio (G_t_ = t_2_/t_1_) for the corresponding durations.

The experiment consisted of two variants of the I(VM) condition. In the first condition, we set d_1_ to 8 degrees and d_2_ to 10 degrees (G_d_=1.25)and in the second condition both d_1_ and d_2_ were 10 degrees (G_d_ =1). In the training sessions, similar to experiment 1 and 2, the stimulus speed was constant throughout the interception path. In the test sessions, unbeknownst to the subjects, immediately after the stimulus entered the occluded segment, its speed was multiplied by 1.25. This manipulation changed G_t_ to 1 and 1/1.25 in the first and second conditions respectively. In other words, in the first condition, G_d_ =1.25 and G_t_ =1, whereas in the second condition G_d_ =1 and G_t_ =1/1.2 5. This manipulation allowed us to tease apart the effect of distance and temporal identity contexts (**Fig. 4a**).

### Analysis

We defined the actual time-to-contact (*TTCa*) as the interval between when the bar passed the central fixation to when it reached the target position. The produced time-to-contact (*TTCp*) was defined as the time from when the bar passed the central fixation to when the subject pressed a key. Subjects that were not sensitive to the range of sample intervals during the training session or had unstable performance were excluded from the study. We considered a subject insensitive if the corresponding *TTCp* distribution for the longest *TTCa* was not significantly different from *TTCp* distribution for the shortest *TTCa* (paired t-test at p=0.05 level). Performance was considered unstable if the first and second order statistics of *TTCp* were different between the first and second halves of the session (paired t-test at p=0.05 level).

### Summary statistics

Following Jazayeri and Shadlen (2010), we characterized each subject’s performance by computing the following summary statistics for *TTCp*:

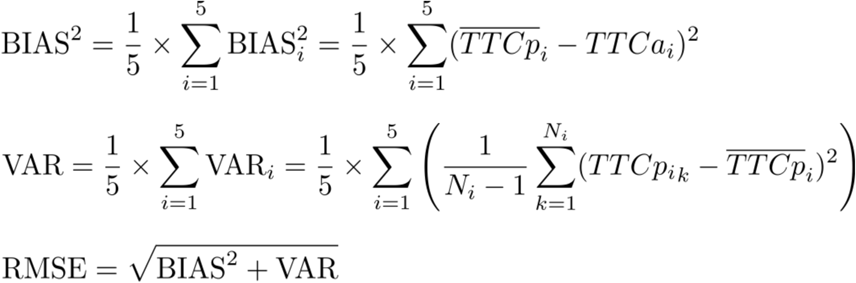

BIAS and VAR represent the average deviations and average variance over the five intervals included in the prior distribution. BIAS_i_ and VAR_i_ represent the mea n deviation and variance of produced times (*TTCp*) for the *i*-th actual interval (*TTCa*) with *N_i_* trials. It follows naturally that the overall root mean squared error (RMSE) is equal to the square root of the sum of BIAS^2^ and VAR. To estimate the mean and variance of summary statistics for individual subject in each condition, we resampled data with replacement and repeated this resampling 100 times

### Effect size

It is known that a relatively large sample size could lead to a smaller p value. Since there were more than 100 trials in each session for each subject, we also measured the strength of difference between conditions for each subject. We used Hedges’ *g^66^*, which is a measure to correct the bias in Cohen’s *d* to quantify the distance between two distribution means. *g* = 0.2 means small effect size, 0.5 means medium effect size, and 0.8 means large effect size.

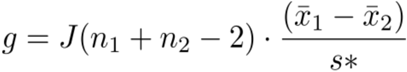

Where

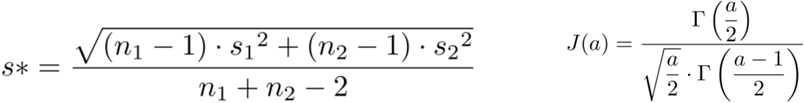

### The Bayesian observer model

We developed a Bayesian observer model (**Fig. 5a**) based on previous work on interval reproduction^24^. We modeled the prior distribution over sample intervals (*t_1_*) based on the ratio of the visible distance (*d_1_*) to the bar’s speed (*v*). To simplify derivations, we modeled the discrete prior distributions used in the experiment as a continuous uniform distribution ranging from the shortest to longest sample interval tests. The shortest and longest intervals were computed in terms of the smallest and largest speeds (*v*^min^ and *v*^max^).

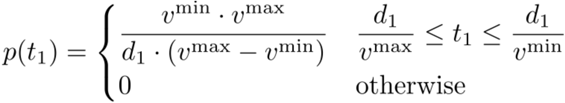

We assumed that subjects made two conditionally independent measurements when the bar was visible, one associated with the speed of the bar, and another associated with duration of the visible period. Following previous work on sensory measurements of time and speed^26^, ^33–36^, we assumed that both measurements were perturbed by scalar Gaussian noise. Specifically, we assumed that the standard deviation of measured speed (*v_m_*) scales with speed (*v*) with constant of proportionality *wmV*, and that the standard deviation of measured elapsed time (*t_m_*) scales with elapsed time (*t_1_*) with constant of proportionality*w_mT_*. The variables*w_mV_* and *w_mT_* represent the Weber fraction for measurement of speed and time, respectively. From the perspective of the observer who makes a measurement *t_m_* and *v_m_*, but does not know *t_1_* and *v*, the problem can be written in terms of the corresponding likelihood functions λ(*t_m_*| *t_1_*) and λ(*v_m_*| *v*):

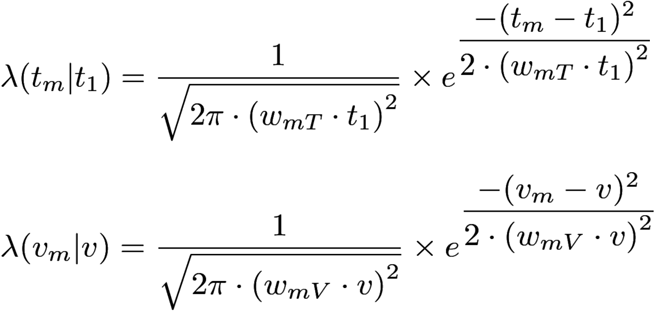

To be able to combine the two likelihoods, we rewrote the likelihood associated with speed in terms of distance and sample interval, as follows:

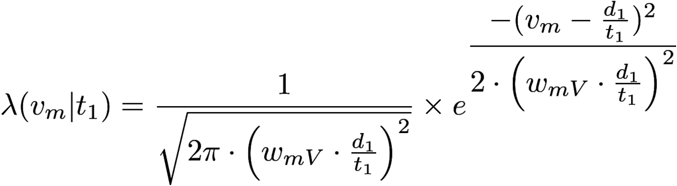

Assuming that the two measurements were conditionally independent, the posterior would be

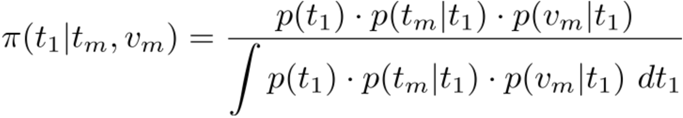

Following previous work^24^, we assumed that subjects’ minimized expected loss using a quadratic loss function, and modeled the inferred duration of the visible period based on the Bayes least-squares (BLS) estimator (i.e., mean of the posterior). We assumed that the estimate was multiplied with a lossless gain (G) to get time-to-contact compute (*TTCe*).

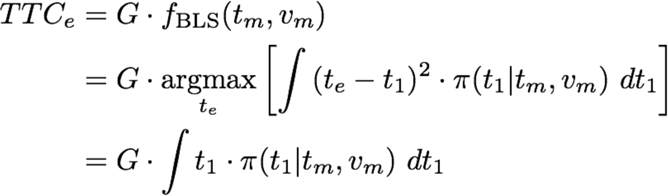

For a uniform prior for sample interval (t1), the time-to-contact estimate (*TTCe*) would be

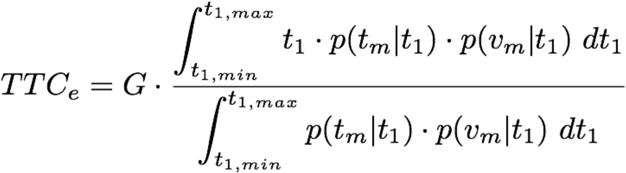

The model was augmented by post-estimation noise to account for motor variability in the produced timeto-contact (*TTCp*). Following previous work^24^, ^33^, ^34^, ^67^, we assumed that the standard deviation of motor noise was proportional to *TTCe*, with constant of proportionality of *w_p_* (Weber fraction for production). We included an offset term (*b_0_*) in the fitting procedure to account for idiosyncratic stimulus- and prior-independent biases observed in responses.

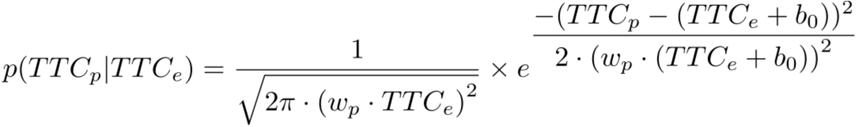

Using chain rule and marginalization of hidden variables, we wrote the the conditional probability of produced time-to-contact (*TTCp*) for a each actual time-to-contact (*TTCa*) as follows:

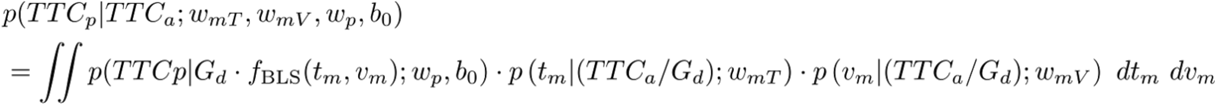

### Fitting procedure

For fitting procedure, we assumed that *TTCp* values associated with any *TTCa* were independent across trials and thus expressed the joint conditional probability of individual *TTCp* values across all the N trials by the product of their individual conditional probabilities.

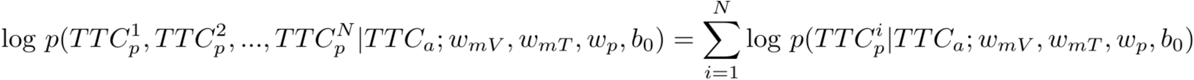

We used fminsearch algorithm to find the model parameters that maximized the likelihood of model parameters across all *TTCa* and *TTCp* values measured psychophysically. Integrals were approximated numerically using the global adaptive quadrature^68^. We repeated the search with different initial values 10 times, and verified that the likelihood functions were stable with respect to initial values.

### Predicting behavior in temporal identity context

Instead of fitting the Bayesian model to each dataset, we asked whether we could fit the model to some conditions and then use parameters of the fitted model to *predict* behavior in other conditions. We aimed to predict behavior in the most important condition where subjects integrated speed with the identity temporal context; i.e., I(VM) with G = 1. We assumed that the noise associated with the measurement of t_1_ is the same in the I(VM) and I(FP) tasks and therefore, used the Bayesian model to the I(FP) task for G = 1 to estimate w_mT_ (**Fig. 5b, left**). We further assumed that the measurement of speed in I(VM) task would be the same across two different gains (G = 1 and G = 0.667), given that the accuracy of speed measurement saturated rapidly (**Supplementary Fig. 2**). We first found w_mT_ for G = 0.667 from I(FP) task, and then used this value to fit a Bayesian model to I(VM) task with G = 0.667 to estimate w_mV_ (**Fig. 5b, right**). Finally, we used the w_mT_ inferred from I(FP) with G = 1 and w_mV_ inferred from I(VM) with G = 0.667 to predict behavior in the I(VM) task with G = 1 in (**Fig. 5c**).

## Acknowledgements

We thank Rossana Chung and Feitong Yang for comments on the manuscript. M.J. is supported by NIH (NINDS-NS078127), the Sloan Foundation, the Klingenstein Foundation, the Simons Foundation, the McKnight Foundation, the Center for Sensorimotor Neural Engineering, and the McGovern Institute.

